# Genome-wide Search for Zelda-like Chromatin Signatures Identifies GAF as a Pioneer Factor in Early Fly Development

**DOI:** 10.1101/104067

**Authors:** Arbel Moshe, Tommy Kaplan

## Abstract

**Motivation:** The protein Zelda was shown to play a key role in early Drosophila development, binding thousands of promoters and enhancers prior to maternal-to-zygotic transition (MZT), and marking them for transcriptional activation. Recently, we showed that Zelda acts through specific chromatin patterns of histone modifications to mark developmental enhancers and active promoters. Intriguingly, some Zelda sites still maintain these chromatin patterns in Drosophila embryos lacking maternal Zelda protein. This suggests that additional Zelda-like pioneer factors may act in early fly embryos.

**Results:** We developed a computational method to analyze and refine the chromatin landscape surrounding early Zelda peaks, using a multi-channel spectral clustering. This allowed us to characterize their chromatin patterns through MZT (mitotic cycles 8-14). Specifically, we focused on H3K4me1, H3K4me3, H3K18ac, H3K27ac, and H3K27me3 and identified three different classes of chromatin signatures, matching “promoters”, “enhancers” and “transiently bound” Zelda peaks.

We then further scanned the genome using these chromatin patterns and identified additional loci - with no Zelda binding - that show similar chromatin patterns, resulting with hundreds of Zelda-independent putative enhancers. These regions were found to be enriched with GAGA factor (GAF, Trl), and are typically located near early developmental zygotic genes. Overall our analysis suggests that GAF, together with Zelda, plays an important role in activating the zygotic genome.

As we show, our computational approach offers an efficient algorithm for characterizing chromatin signatures around some loci of interest, and allows a genome-wide identification of additional loci with similar chromatin patterns.

## 1 Introduction

The process of transcription is vital to all living organisms, and is tightly regulated by multiple mechanisms, including the packaging of DNA into chromatin. In eukaryotic cells, The DNA is wrapped around nucleosomes to form chromatin. This packaging is used differentially to control in what conditions and cell types a gene is more accessible - and active - and in which conditions it is tightly packed and silenced. This packaging of DNA was shown to be mediated by various mechanisms, including the deposition of covalent modifications (e.g. acetylation, methylation, phosphorylation or ubiquitylation) at different residues of the core histone proteins, that are assembled into a nucleosome (Kouzarides 2007). These histone modifications influence various processes along the DNA. For example, H3K4me1 and H3K27ac (namely, mono-methylation of Lysine 4 or acetylation of Lysine 27 in histone H3) are found at nucleosomes carrying active regulatory regions, while H3K27me3 is known to be a repressive mark (Liu et al. 2005, Heintzman et al. 2007, Rada-Iglesias et al. 2011).

The packaging of DNA into chromatin, including nucleosome positioning and their histone modifications, are fundamental to the proper activity of regulatory regions, by controlling which regions of the genome are accessible for protein binding and which are not (Kaplan et al. 2011, Thomas et al. 2011, Li et al. 2014). Of specific interest are *promoter* regions, located near the transcription start site (TSS) of genes; and *enhancers*, that regulate gene expression from afar, often up to 1Mb away from their target genes. In addition, chromatin marks and structural proteins allow distal enhancers to fold in 3D into close spatial proximity to their target genes (Schübeler 2007, Zaret and Carroll 2011, Smallwood and Ren 2013).

Comparison of tissue- and condition-specific chromatin data highlight the tight regulation of gene expression by chromatin, determining which genomic regions are active and which are not, and as a consequence which genes will be transcribed. This raises the question of causality. Who regulates packaging? Or how does the genome get packed initially in the proper architecture, e.g. to drive early developmental expression?

**Figure 1.**
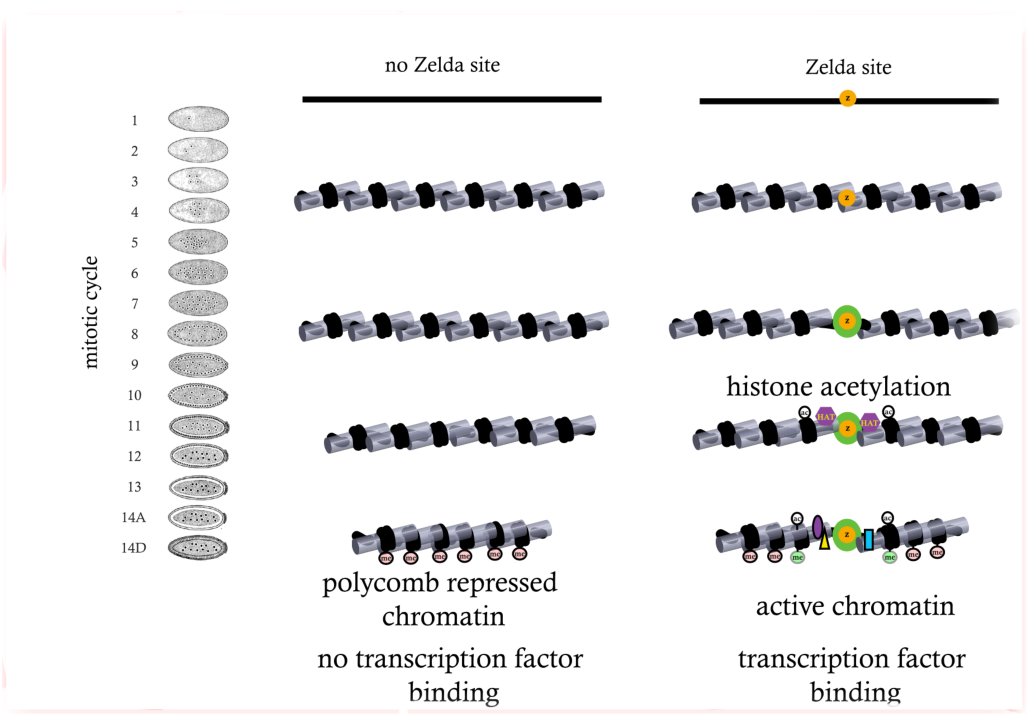
The role of Zelda during zygotic genome activation. Zelda binds to regulatory regions in pre-MZT embryos as early as mitotic cycle 2. This leads to histone acetylation and nucleosome remodeling around ZLD binding sites, which facilitates binding by other transcription factors and deposition of active histone marks. At the same time, Zelda binding prevents deposition of H3K27me3 marks and formation of repressive chromatin structure. (adapted from Li et al., 2014)

In practically all animals, early developmental stages begin with maternal proteins and RNA that control the first hours in the fertilized egg. At this stage, these proteins control the first wave of zygotic expression and direct the first mitotic divisions (Tadros and Lipshitz 2009). After that the embryo undergoes a process called Maternal-to-Zygotic Transition (MZT), in which the zygotic genome is activated and takes control of mRNA and protein production. Finally, maternal mRNA and proteins are degraded.

In the fruit fly *Drosophila melanogaster*, embryonic development is characterized by a series of 13 rapid replication cycles, occurring during the first two hours after fertilization. The division of cells slows at the 14^th^ mitotic cycle, and zygotic transcription initiates. This marks the end of the *Drosophila melanogaster* Maternal-to-Zygotic Transition. This process is crucial for the normal development of the embryo and is tightly controlled, both in time and space, by the gradual activation of a cascade of transcription factors (Li et al. 2008, Macarthur et al. 2009). This required multiple molecular mechanisms that include chromatin, nucleosomes, DNA accessibility, steric hindrance between DNA binding proteins and more.

Previous research showed that many of the early-transcribed genes in *Drosophila* embryos contain a specific DNA motif of 7bp, CAGGTAG, that occurs within their regulatory regions, including both promoters and enhancers (ten Bosch et al. 2006, Bradley et al. 2010, Harrison et al. 2011). This motif was later identified to be the binding site of the zinc-finger transcription factor Zelda (vielfaltig, vfl) (Liang et al. 2008). Following studies, by us and others, showed that Zelda is present in the embryonic nucleus as early as mitotic cycle 2 and binds thousands of genomic loci, including the promoters and enhancers of thousands of early developmental genes (Figure 1) (Li et al. 2008, Liang et al. 2008, Harrison et al. 2011, Nien et al. 2011).

Computational and experimental studies, by us and others, suggested that Zelda acts as a Pioneer Factor, binding mostly inaccessible DNA regions and making the chromatin accessible for other transcription factors to bind, thus marking thousands of genes and regulatory regions for activation which drives the first transcriptional program of the developing embryo (Harrison et al. 2011, Zaret and Carroll 2011, Satija and Bradley 2012).

Indeed, experimental studies showed that early Zelda binding is a predictor for open chromatin and transcription factor binding at mitotic cycle 14 (Liang et al. 2008, Macarthur et al. 2009, Bradley et al. 2010, Harrison et al. 2011, Schulz et al. 2015). The entire set of molecular mechanisms by which Zelda functions to access the genome and mark it for activation is yet to be discovered, as is the role of additional proteins in this crucial stage in embryonic gene expression.

Several studies mapped the chromatin landscape in *Drosophila melanogaster*, most of which in cells or during very broad temporal windows (Filion et al. 2010, modENCODE Consortium et al. 2010, Kharchenko et al. 2011, Nègre et al. 2011). Of particular interest are few studies, in which early and manually staged *Drosophila* embryos were used to portray Zelda binding locations and multiple histone marks (including H3K27ac, H3K4me1, H3K4me3, H327me3, and H3K36me3) at several time points throughout MZT and early embryonic development (Harrison et al. 2011, Li et al. 2014). As was shown, early Zelda binding often results in open chromatin and characteristic histone marks, including H3K27ac, H3K4me1, and H3K27me3 peaks (Liang et al. 2008, Sun et al. 2015).

Intriguingly, the causal role of Zelda in proper establishment of chromatin domains and normal gene expression was also examined. Embryos lacking zygotic Zelda expression, as well as embryos with mutations in specific Zelda binding sites (near early genes) showed developmental abnormalities (ten Bosch et al. 2006, Liang et al. 2008). Embryos lacking maternal Zelda expression (*zld*^*M-*^) showed reduced accessibility for many, but not all, distal enhancers regulating early fly development, while maintaining near-normal promoter accessibility (Li et al. 2014, Schulz et al. 2015).

The latter results suggest that perhaps, in addition to Zelda, there might be another protein that marks early developmental genes for activation. To answer this question, we revisited the chromatin patterns surrounding Zelda binding sites, and developed a computational statistical model to characterize histone modifications and their dynamics. At a second stage, we used this computational model to scan the genome and identify additional genomic loci that show similar chromatin patterns.

## 2 Results

To identify additional Zelda-like regulators that act as pioneer factors during early *Drosophila* development, we begin by characterizing the chromatin landscape induced around early Zelda sites.

### 2.1. Zelda peaks vary in their chromatin signatures

We have previously shown that early Zelda peaks, identified via ZLD ChIP-seq in hand-sorted fly embryos from mitotic cycle 8, are associated with open chromatin regions and transcription factors binding later on, towards the end of the Maternal-to-Zygotic Transition (mitotic cycle 14) (Figure 1) (Li et al. 2008, Macarthur et al. 2009, Harrison et al. 2011). We begin by portraying the chromatin landscape surrounding these early Zelda sites. As shown in Figure 2A, many Zelda peaks are accompanied with strong H3K4 mono-methylation, and depletion of H3K27me3 and H3K36me3 (Harrison et al. 2011).

**Figure 2.**
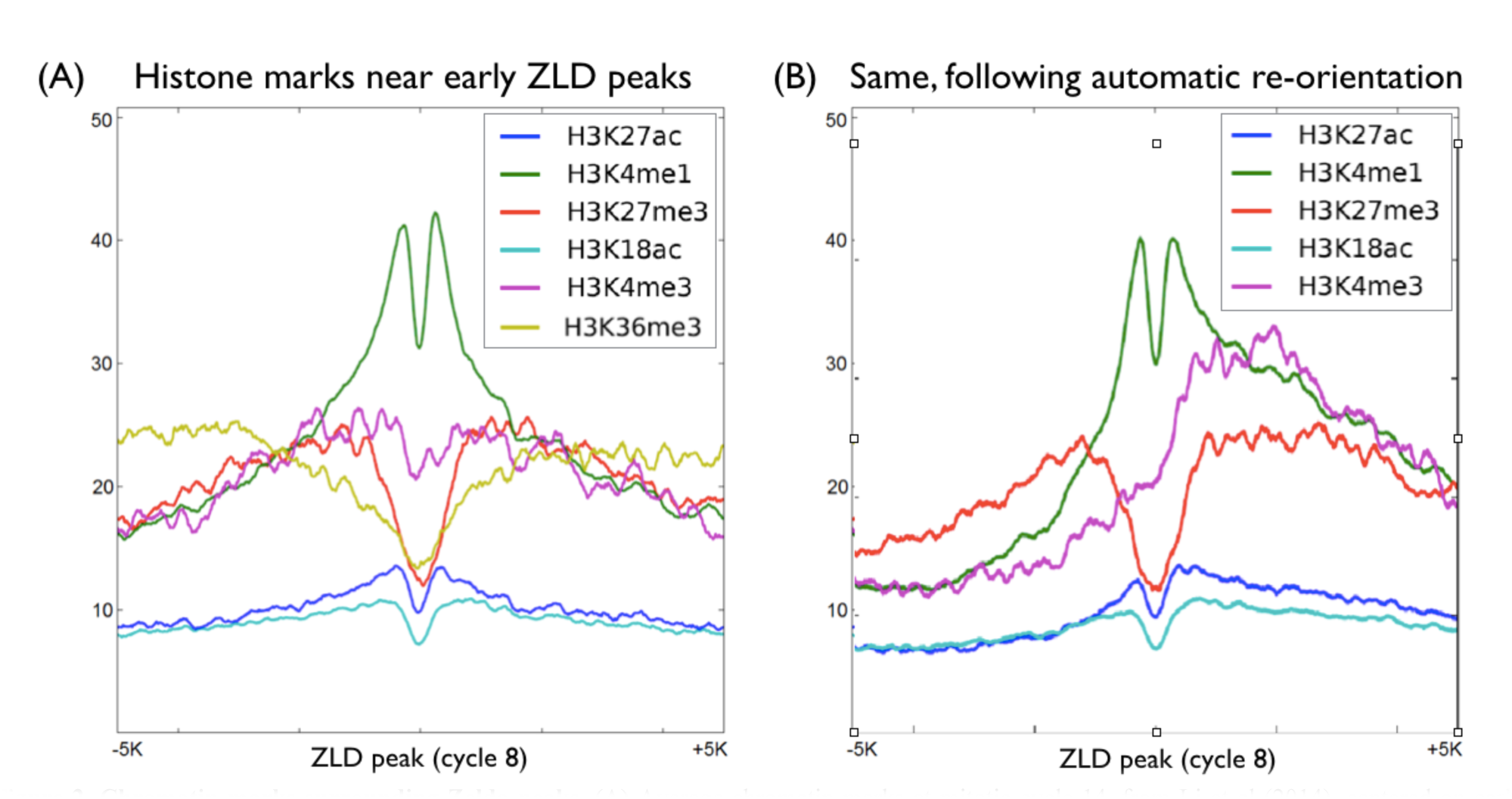
Chromatin marks surrounding Zelda peaks. **(A)** Average chromatin marks at mitotic cycle 14, from Li et al (2014), centered on early Zelda peaks (cycle 8). Clearly visible are relative enrichment near Zelda site for the enhancer mark H3K4me1 (green), and depletion of the repressive H3K27me3 (red). **(B)** Same, following peak re-orientation.

However, a close examination of some strong early Zelda sites (Figure 3) suggests that there is more than one typical chromatin signature. Specifically, shown are two Zelda peaks, one at an intron of the *schnurri* gene (*shn*, Fig. 3A), and one at the promoter of *bitesize* (*btsz*, Fig. 3B). The former shows a ZLD peak surrounded by H3K4me1-marked nucleosomes and flanked by H3K27 tri-methylation. The latter shows the promoter Zelda peak is surrounded by H3K4me1 nucleosomes, as well as H3K4me3 on one side of the ZLD peak. Intriguingly, as shown in Fig. 3C, some early Zelda peaks seem to show no chromatin pattern whatsoever (by cycle 14), and are practically indistinguishable from their surroundings.

**Figure 3.**
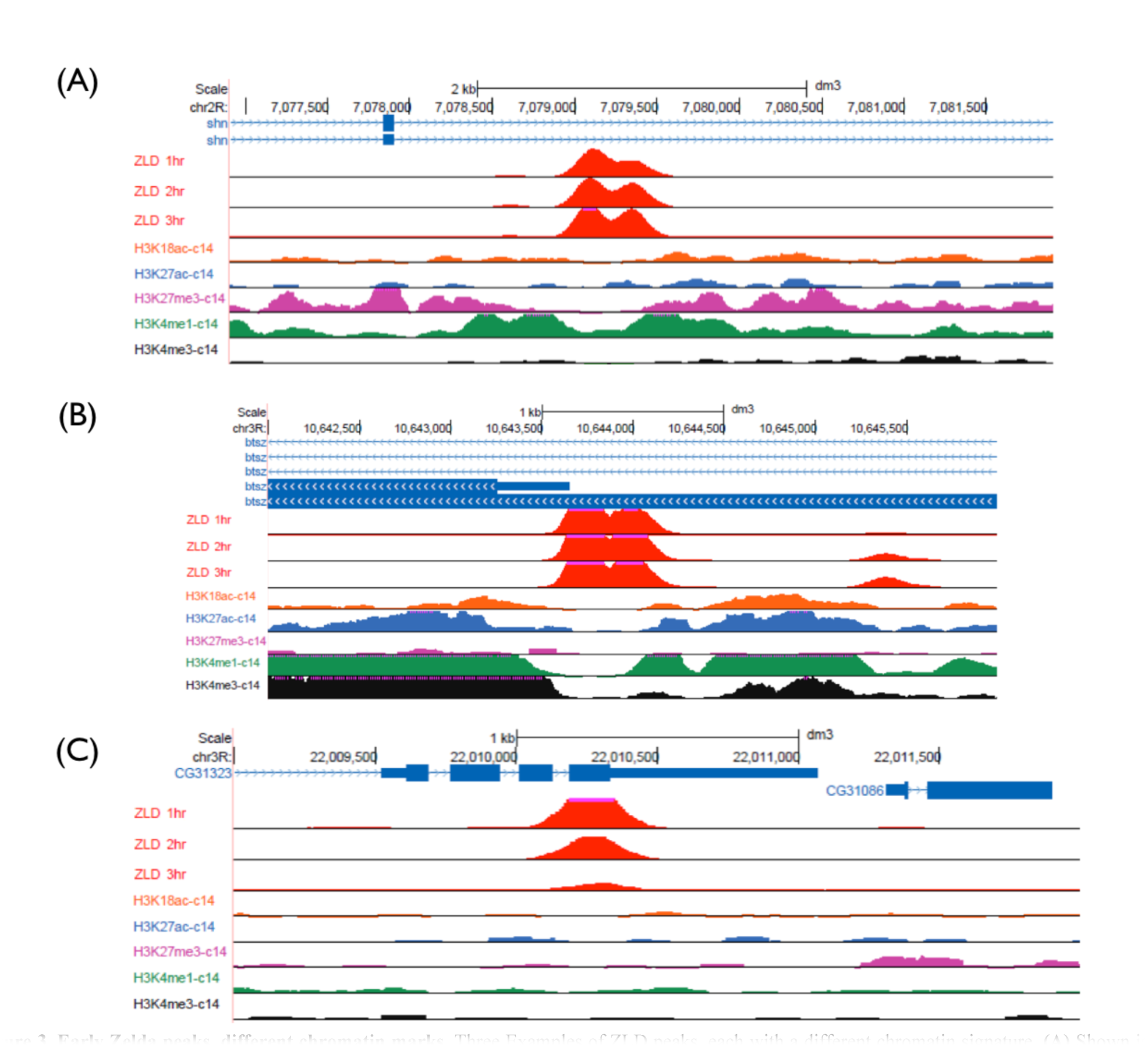
Early Zelda peaks, different chromatin marks. Three Examples of ZLD peaks, each with a different chromatin signature. **(A)** Shown is a distal enhancer of the *schnurri* gene (*shn*), where Zelda binds and an intronic enhancer (also bound by multiple A-P transcription factors, not shown), is flanked by H3K4me1 nucleosomes (green) within an H3K27me3 domain (purple). **(B)** Shown is the promoter of the *bitesize* gene (*btsz*), highlighted with the typical promoter modification (H3K4me3, black) and an asymmetric H3K4me1 domain (green), with flanking H3K27ac and H3K18ac flanking nucleosomes (blue, orange). **(C)** Shown is a ZLD peak overlapping the coding region of an inactive gene – zygotic expression is smaller than 0.2 FPKM in mitotic cycles 10-14 (Lott et al. 2011). Zelda seems to be transiently bound, with very weak binding by the end of mitotic cycle 14 (Harrison et al. 2011). No histone modification are observed along the peak’s neighboring regions.

We therefore hypothesized that different Zelda peaks could have different histone modification patterns, which may be obscured when only considering the average, common pattern, near Zelda peaks.

### 2.2. Chromatin-based re-orientation of early Zelda peaks

As a first step towards a more descriptive and accurate characterization of chromatin near Zelda sites, we first wanted to check which histone modifications are symmetric on both sides of the Zelda peaks, and which are not. For this, we performed an Expectation Maximization (EM)-like iterative orientation method based on histone modification data, and automatically inferred the orientation of each Zelda peak, while iteratively improving the typical chromatin “signature” surrounding Zelda peaks (Dempster et al. 1977).

We decided to choose a fully unsupervised chromatin-based reorientation, rather than relying on gene directionality, as we believe that at least some of the modifications are “one-sided”, and therefore would contribute to inferring the orientation of most Zelda peaks. Indeed, while most of the modifications maintain a symmetrical formation, both H3K4me3 (promoter) and H3K36me3 (gene body) show strong asymmetric bias, consistent with the orientation of the underlying gene (Figure 2B).

### 2.3. Chromatin-based clustering of Zelda peaks

To test whether there are different types of Zelda peaks, each with a different combination of histone modifications patterns, we turned to develop a computational model that will allow us to characterize the chromatin landscape and temporal dynamics at each early Zelda peak. Such an approach would not only shed light on the different functional roles of Zelda peaks, but would also allow us to scan the genome using a refined model, and identify additional loci, independent of ZLD binding, that undergo similar chromatin dynamics during MZT.

As shown in Figures 2 and 3, it is not enough to only consider the maximal signal of each histone modification, as the modifications we look at are not characterized by a single peak, but rather show a unique spatial signature, which is often asymmetric. We therefore want a specialized method that considers both the pattern of each histone modification and the overall combination of different modifications and their height.

For this, we decided to examine a 10Kb window surrounding each Zelda peak, and apply a spectral clustering algorithm that will consider all histone modifications simultaneously. We begin by describing a distance function between the chromatin signature of two loci that would capture both the “landscape shape” of each modification, its overall magnitude, and the combination of multiple modifications.

An advantage of spectral clustering is the fact that the algorithm relies on the adjacency (or connectivity) between objects within each cluster, rather than their spatial shape.

Our initial chromatin ChIP-seq data consisted of nine histone modifications (H3K4ac, H3K4me1, H3K4me3, H3K9ac, H3K18ac, H3K27ac, H3K27me3, H3K36me3, and H3K5ac) in four time points throughput MZT (mitotic cycles 8, 11, 13, and 14) (Li et al. 2014). We therefore selected the top 2,000 early (cycle 8) Zelda peaks, and for each considered a surrounding window of 10Kb for five regulatory histone modifications (H3K4me1, H3K4me3, H3K27ac, H3K27me3, and H3K18ac), as measured in mitotic cycle 13 and 14. While H3K36me3 showed some depletion near Zelda sites (Figure 2A, gold), we decided not to use it for the clustering as it is mostly found along gene bodies, and did not show additional information. The idea here was not to obtain a definitive metric for chromatin, but rather to identify the regulation-specific chromatin landscapes near Zelda peaks.

This resulted in representing each genomic locus **
*i*
** by ten vectors 
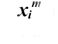
 one for each of the five histone modifications at each of the two time points (marked by ***m***). We have down-sampled the chromatin data to 10bp resolution, resulting in vectors of length 1,000. To calculate the distance 
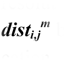
 between the two vectors representing the landscape of the modification ***m*** surrounding two loci ***i*** and ***j***, we first used the root mean squared difference (RMSD, see Equation 1, Methods) between the two vectors 
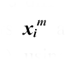
 and 
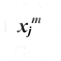
 then converted these into weights (or adjacency scores), using a Gaussian kernel with a modification-specific parameter ***σ***_***m***_ (Equation 2, Methods). This parameter sets the standard deviation parameter of the Gaussian kernel, thus normalizing different histone modifications and assigning each a similar importance. We have arbitrarily set ***σ***_***m***_ for each modification/time point to equal to the 10^th^ percentile in the distribution of pairwise distances for that specific modification (see Methods). Finally, we summed each of the ten adjacency matrices (Equation 3), and applied standard (symmetrically normalized) spectral clustering, followed by k-means (Ng et al. 2001). The value for ***k*** (i.e., number of clusters) was chosen using the eigengap heuristic (Methods). Overall, the clustering process resulted with three main clusters, each with a unique combinatorial chromatin signature (Figure 4).

**Figure 4.**
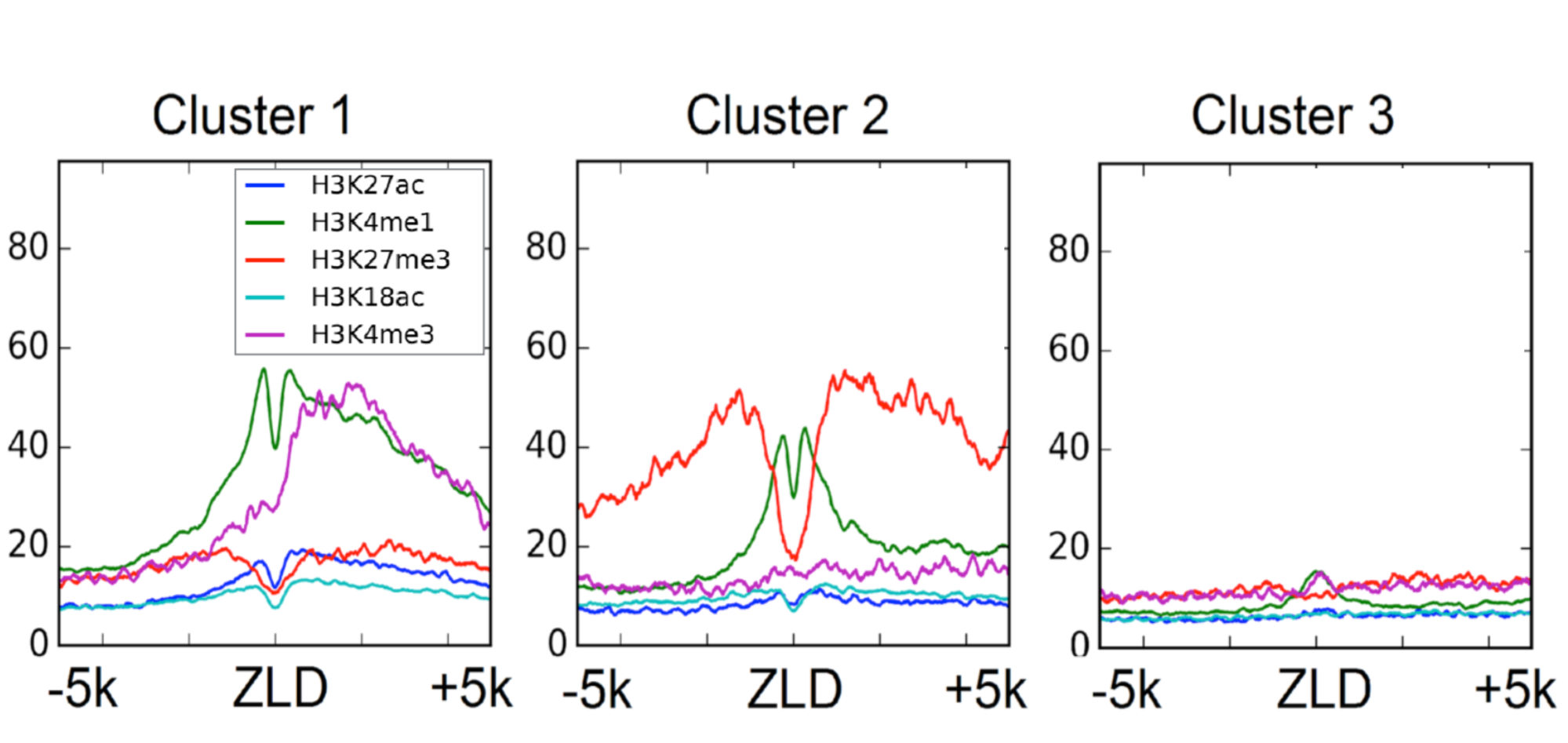
Three typical chromatin signatures found among top 2,000 early Zelda sites. Cluster 1 (left) is composed of 1,032 early Zelda peaks, with an average chromatin signal of asymmetric H3K4me3 with H3K4me1 mark (promoter like signature). Cluster 2 include 465 early Zelda peaks, and shows symmetric signature of two flanking H3K4me1 nucleosomes, within a larger H3K27me3 region. Cluster 3 (right) includes 503 Zelda peaks, and is characterized by almost no modifications.

As reported by Li et al. (2014), most of these modification show little change between cycles 13 and 14, and we therefore only plot the former here. The first cluster (Figure 4, left) is composed of 1,032 early Zelda peaks (of the initial 2,000 peaks analyzed), and shows enrichment for H3K4me1 (green), on either side of early Zelda peaks, and H3K4me3 (purple), enriched only on one side of Zelda peaks.

The second cluster (Figure 4, center, 465 Zelda peaks) shows narrow symmetric H3K4me1 peaks on either sides of the Zelda peak, with wider H3K27me3 domains further away. Intriguingly, while this pattern resembles the chromatin signature of “poised enhancers”, described by Rada-Iglesias et al. (2011) in human embryonic stem cells (hESCs), a closer examination suggest that most of these genomic regions are actually active enhancer regions (see below).

Finally, the third cluster (Figure 4, right, 503 Zelda peaks) showed almost no enriched chromatin marks, suggesting that these are transient Zelda binding sites that are bound early and then vacated with no apparent biological effect.

### 2.4. Functional characterization of three Zelda clusters

As we showed, spectral clustering of early Zelda peaks resulted in three clusters, each displaying a unique combination of histone modifications. We next turned to examine whether these clusters demonstrate different functional parameters.

First, we wanted to test if the detected differences in chromatin packaging affect transcription factor binding. For this, we analyzed genome-wide binding data for 21 early development transcription factor, as measured after the completion of the Maternal-to-Zygotic transition at mitotic cycle 14 (Macarthur et al. 2009). For each of the 2,000 early Zelda peaks that were clustered, we calculated the number of different transcription factors that are bound at the same locus. As shown in Figure 5A, clusters 1 and 2 show an average of >7 bound transcription factors (at cycle 14), compared to cluster 3 where only <2 bound transcription factor were found on average.

**Figure 5.**
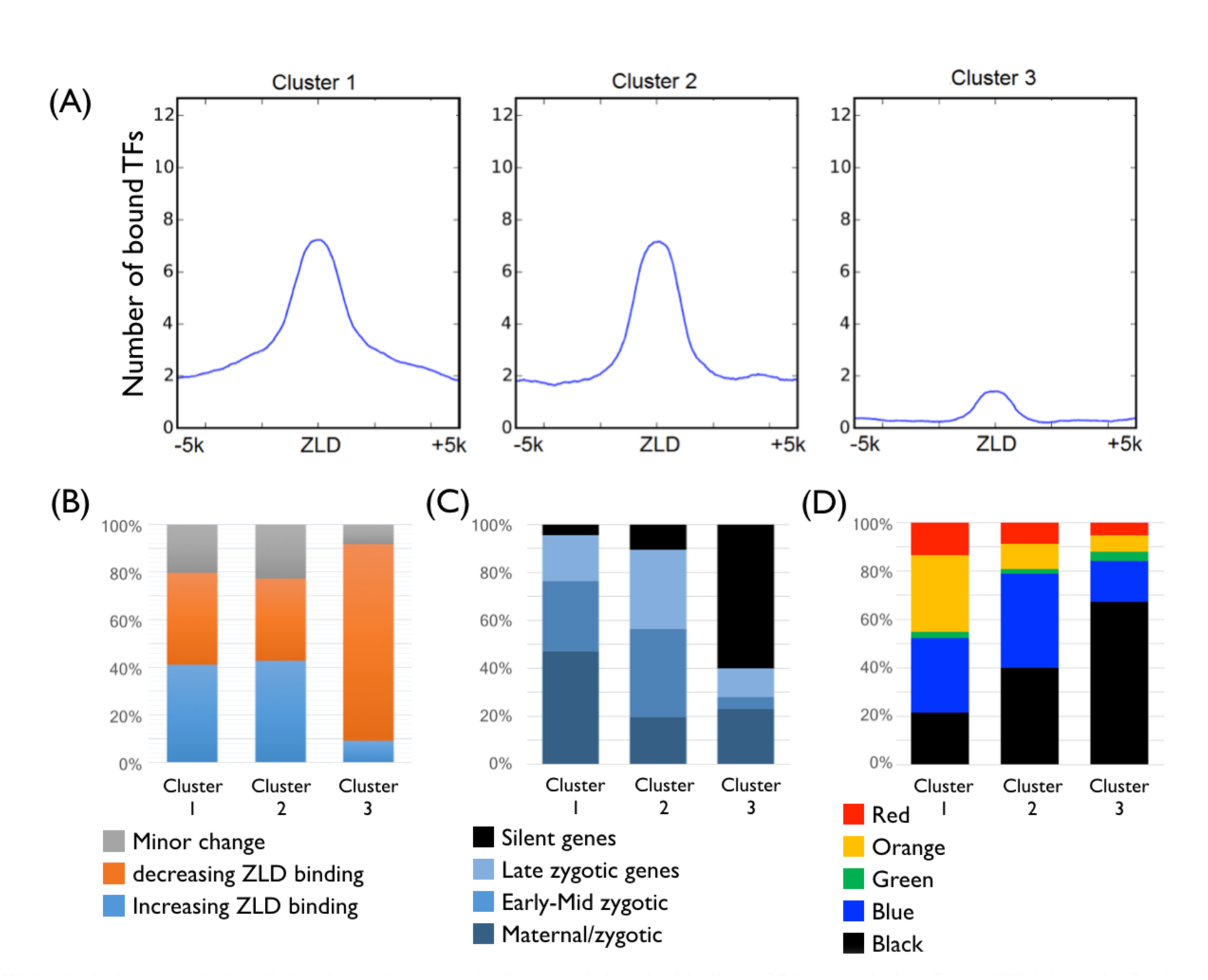
Biological characteristics of the three clusters. **(A)** By examining the binding of 21 transcription factor (TFs) in early fly development, we calculated the average number of TFs bound for each locus with each cluster (Macarthur et al. 2009). Early Zelda peaks in clusters 1 and 2 seem to be bound (by the end of mitotic cycle 14) by ~7 TFs, compared to <2 TFs for Zelda peaks in cluster 3. **(B)** Analysis of temporal dynamics in Zelda binding (Harrison et al. 2011). In clusters 1 (left) and 2 (center), about 40% of the ZLD peaks are increasing, 40% decreasing, and 20% show minor change in peak height. In contrast, more than 80% of cluster 3 regions (right) show reduced ZLD binding. **(C)** By associating each enhancer-like locus to the nearest TSS, we show that cluster 1 loci are mostly associated with maternally deposited genes that are also transcribed in early developmental stages (“Maternal/ zygotic”, e.g. house-keeping genes). Cluster 2 is more associated with zygotic genes, while cluster is mostly not expressed (Lott et al. 2011). **(D)** Different chromatin patterns as annotated using HMM with 53 chromatin proteins in Drosophila Kc167 cells (Filion et al. 2010).

This raises the hypothesis that while most early Zelda sites become accessible during MZT and facilitate the binding of regulatory proteins, perhaps the sites captured by cluster 3 do not.

For this, we turned to examine the dynamics of Zelda binding itself, as measured by ChIP-seq in three time-points throughout MZT (Harrison et al. 2011). As shown in Figure 5B, similar number of early Zelda peaks exhibit increased- or decreased ChIP signal during MZT. Conversely, almost all peaks in cluster 3 showed a decrease in Zelda binding, consistently with the observed reduction in transcription factor binding.

To test the functional effect of these three groups in regulating gene expression, we associated each Zelda peak with the nearest gene. Indeed, we observed differences in the distance distribution to the nearest TSS. While most cluster 1 peaks were directly located at gene promoters (hence the asymmetric H3K4me3 signature shown in Figure 4, left), peaks from cluster 2 were further away from the TSS (typically 2.5- 3Kb). Considering the chromatin signature (H3K4me1 and H3K27me3) and the high number of TF bound at these loci, it is not unlikely that most of the distal regions (in cluster 2) act as regulatory enhancers.

Following Harrison et al. (2011), where we used single-embryo RNA-seq data from mitotic cycles 10-14 (Lott et al. 2011) to classify early developmental genes into temporal groups, we have now compared these annotations to the genes associated with each of the peaks in the three clusters. As shown in Figure 5C, peaks in cluster 1 were mostly associated with maternal and zygotic genes. Zelda peaks from cluster 2 (i.e. the putative enhancer-like group) were even more enriched with early, strictly-zygotic genes. Cluster 3 was dramatically enriched with silenced and housekeeping genes (expressed both maternally and zygotically).

Finally, we compared the three clusters to HMM-based annotations of the fly genome into five chromatin types, based on the binding patterns of 53 chromatin proteins in Drosophila Kc167 cells (Filion et al. 2010). Indeed, as shown in Figure 5D, the three classes show district patterns, with cluster 1 mostly enriched for the active genes (yellow, red), as well as genomic regions that are packed in polycomb (blue) or repressed (black) in Kc167 cells.

### 2.5. *de novo* chromatin-based identification of Zelda-like loci

We next turned to use the three chromatin signatures obtained using the spectral clustering algorithm, and identify additional loci showing similar chromatin patterns. While Zelda plays a major role in shaping the chromatin landscape and accessibility during the MZT, marking them for activation and enabling the binding of regulatory proteins, many early *Drosophila* genes are still expressed in Zelda maternal mutants embryos (Li et al. 2014, Schulz et al. 2015). This suggests that there might be some other mechanisms (besides Zelda) that may also lead to similar chromatin signatures of active and regulatory chromatin.

For this, we cross-correlated the genome with the chromatin patterns of the promoter-like (cluster 1) and enhancer-like (cluster 2) signatures (Figure 4). We used the same set of histone modification marks as used for clustering (H3K27ac, H3K27me3, H3K4me1, H3K4me3, and H3K18ac) at cycle 14, with the same 10Kb windowing. Initially, we separately scanned the genome with each of the histone modification patterns and computed the average correlation for all modifications. This naive analysis failed to obtain satisfying results, as the ratio between the different histone modifications was not preserved. As a result, many loci showed high correlation to each of query patterns, but their combined chromatin signature was skewed. Next, we tried to search for all modifications simultaneously by concatenating the histone modifications into one long vector, then cross-correlating against the genome. These results were still rather disappointing, since correlation by itself does not account for the overall magnitude of the histone modification pattern, but only the overall shape.

To solve this issue, we added a regularization term that penalized the correlation coefficient obtained for each genomic locus (Equation 7). This factor was estimated based on the ChIP height distribution for each histone modification among the original Zelda peaks, and reflected the empirical prior probability of obtaining a peak of certain height at positive samples (Methods).

We then scanned the *Drosophila melanogaster* genome, and extracted genomic loci with high local score. As shown in Figure 6, our de novo chromatin-based identification of Zelda-like loci retrieved the majority of bona fide Zelda peaks in clusters 1 and 2. When considering the top 2,500 hits identified by the promoter-like (cluster 1) signature, we identified over 75% of the original Zelda peaks in that cluster, with the addition of ~400 (15%) regions with similar chromatin patterns (Figure 6A). Similarly, by scanning the genome with the enhancer-like (cluster 2) pattern, we managed to locate more than 80% of original early Zelda peaks, with an addition of ~200 (7%) new loci (Figure 6B). When testing for TF binding (as done before, Figure 5B) we observed an average of 5.2 bound factors among the novel enhancer-like regions (compared to ~7 TFs found at the top Zelda-bound regions and <2 factors bound at random regions).

**Figure 6.**
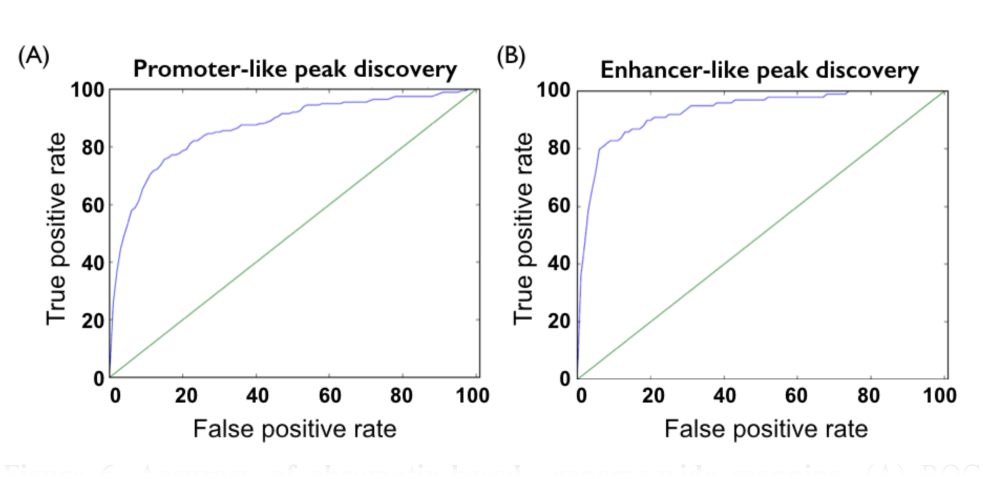
Accuracy of chromatin-based genome-wide scanning. **(A)** ROC curve of the promoter-like chromatin model. Y-axis corresponds to percent of early Zelda peaks (from cluster 1) retrieved by a genome-wide scan for cluster 1 chromatin signatures. X-axis denotes the percent of newly discovered (non-ZLD bound) regions with similar chromatin signature. For example, we retrieved over 75% of the original Zelda peaks, with an addition of ~400 (15%) regions with similar chromatin patterns (denoted as False positives here). **(B)** Similar plot for enhancer-like regions. Here, it is possible to retrieve 80% of the original Zelda peaks, with an addition of ~200 (7%) new loci

### 2.6. Motif Analysis of discovered putative regulatory sites

To further examine the functional role of the newly discovered enhancer-like regions, we ran a *de novo* motif analysis on the top 1,600 enhancer-like peaks, using HOMER (Heinz et al. 2010). Our motif analysis, based On 100bp, 250bp or 500bp long sequences, all identified three major motifs *de novo* (Figure 7).

**Figure 7.**
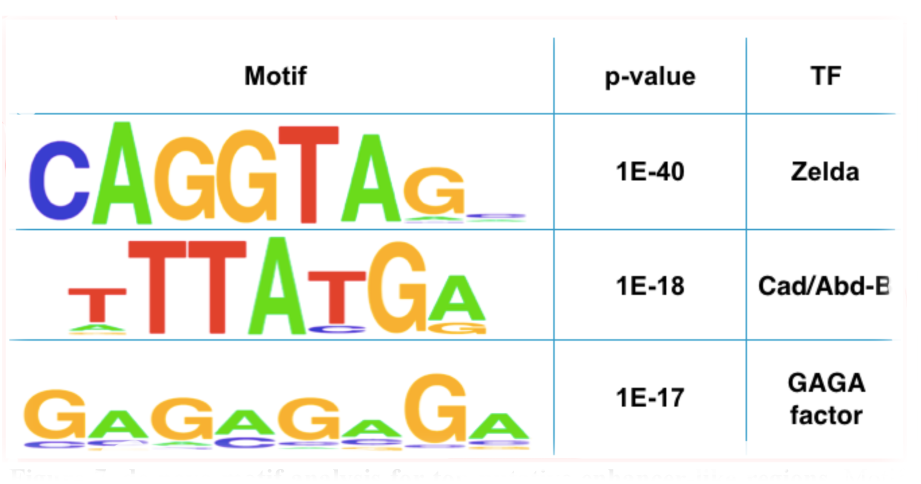
de novo motif analysis for top putative enhancer-like regions. Motif analysis of top 1,600 putative enhancer-like regions using HOMER (Heinz et al. 2010). Using 100bp windows, we identified the CAGGTAG motif (Zelda binding site, p<1e-40), a second motif (p<1e-18, possible related to Caudal or Abd-B), and poly(GA), bound by GAGA factor (GAF, Trl).

The most significant motif identified (1e-40) is CAGGTAG, the known recognition element of Zelda (ten Bosch et al. 2006, Liang et al. 2008, Bradley et al. 2010, Harrison et al. 2011). Not surprising, considering the enrichment of Zelda sites within our set of chromatin-based enhancer-like loci. The second motif identified by HOMER, with a p-value of 1e-18 is TTATGA, and is similar to the known recognition site of two *Drosophila* proteins, Caudal (Cad) and Abdominal-B (Abd-B). The latter shows very low expression levels, and only towards the end of cycle 14 (Lott et al. 2011), while the former is a key transcription factor that is expressed in a gradient towards the posterior end of early *Drosophila* embryos and acts as a morphogen for abdomen/tail formation. Motif 3, with a p-value of 1e-17, is a poly(GA)-rich motif, and is similar to the known recognition site of the protein GAGA factor (GAF, also referred to as Trithorax-like, or Trl).

To determine whether Caudal or GAGA factor (GAF) may act as pioneer factors independently of Zelda, we examined ChIP-seq and ChIP-chip data for Caudal and GAF, and calculated their overlap with Zelda peaks (Macarthur et al. 2009, Harrison et al. 2011, Nègre et al. 2011). Overall, almost all Caudal peaks (94%) seem to overlap with Zelda binding, compared to only half (56%) of GAF peaks. We therefore continued to study the role of GAF as a pioneer factor.

### 2.7. GAGA factor (GAF) acts as a Zelda-like pioneer factor

As our motif analysis showed, the GAGA motif (to which GAF binds) is enriched among the putative enhancer-like loci, with about half of GAF sites showing no Zelda binding. To test how prevalent GAF binding is at those enhancer-like loci, we also analyzed GAF ChIP data from hours 0-8 of *Drosophila melanogaster* development (Nègre et al. 2011). Indeed, the average ChIP strength for the predicted enhancer-like loci is more than three-fold higher, compared to flanking regions (77 vs. 22). It should be noted that similarly to the motif analysis where only ~15% of the enhancer-like regions contained a GAGA motif, analysis of ChIP data identified GAF binding in about 20% of these regions (compared to over 75% for Zelda binding).

Finally, we turned to directly test the ability of GAF to act as a pioneer factor with or without Zelda. Recently, we measured DNA accessibility via formaldehyde-assisted isolation of regulatory elements (FAIRE) experiments (Schulz et al. 2015). The data consists of FAIRE throughout 2-3 hours of development in wild-type embryos, as well as embryos maternally depleted for Zelda (*zld*^*M-*^).

To test the hypothesis that GAF and Zelda could both act as a pioneer factors, we decided to analyze ZLD and GAF binding to every putative enhancer-like region, and to compare these data with their overall DNA accessibility data (as measured by FAIRE) in WT and in *zld*^*M-*^ mutants. For ZLD, we expected to see a positive correlation between ZLD binding and DNA accessibility (i.e. regions with strong ZLD binding are more likely to be accessible) in WT, but not in *zld*^*M-*^ mutants. In addition, we expect to see that enhancers with strong GAF binding and weak Zelda binding are affected by the deletion of *zld*, namely to show strong accessibility in both WT and *zld*^*M-*^ mutants FAIRE data.

Due to the relatively low number of data points (2,500 enhancer-like regions divided into 36 groups according to their ZLD and GAF ChIP signals), we only show the average FAIRE accessibility for each group. As Figure 8A shows for the WT embryos, an increase in both ZLD binding (moving upwards) or GAF binding (moving to the right), result in increased DNA accessibility (darker shade of blue). Conversely, the accessibility matrix for *zld*^*M-*^ mutants is less sensitive to ZLD levels, especially on the left hand side, were GAF levels are very low, yet it is almost unaffected by the deletion of zld among the strong GAF sites (Figure 8B, right hand side). These data suggest that GAGA factor may act as a pioneer factor and contribute to the accessibility and chromatin landscape of DNA regions, much like Zelda (yet for fewer genomic loci), independently of Zelda.

**Figure 8.**
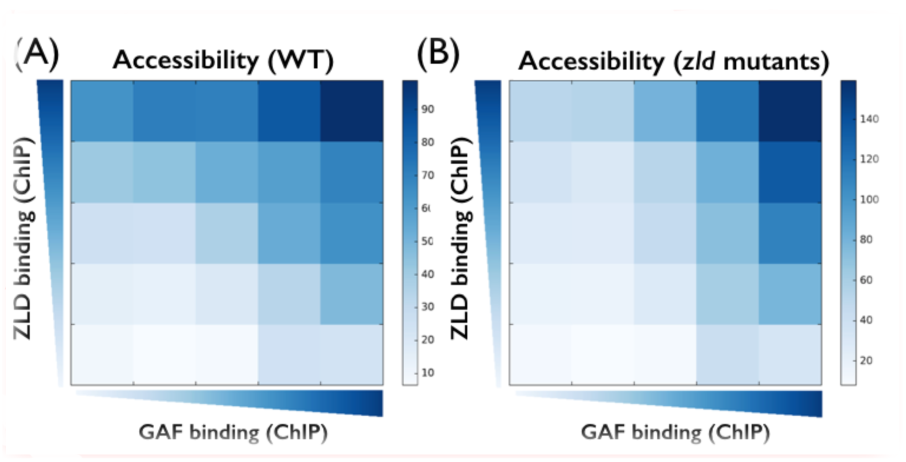
DNA accessibility in WT and *zld* mutants. (A) Shown is the average FAIRE signal (DNA accessibility) in WT embryos for 2,500 enhancer-like loci that are binned based on their ZLD (Y-axis) and GAF (X-axis) ChIP binding. (B) Same as A, using FAIRE data from maternal *zld* mutants (Schulz et al. 2015). Comparison of the two matrices show that genomic regions with no GAF binding (in WT) are mostly affected by deletion of *zld* (left hand side of both matrices), while regions bound by GAF (right hand side) still show high DNA accessibility, suggesting that GAF could ac as a pioneer factor in early fly development, together or independently of Zelda.

## 3 Methods

### 3.1. Multi-channel chromatin representation of ZLD sites

To characterize the chromatin signature around Zelda sites, we considered the top 2,000 Zelda peaks in mitotic cycle 8 (Harrison et al. 2011), and looked at five histone modifications (H3K27ac, H3K27me3, H3K4me1, H3K4me3 and H3K18ac) at two time points (mitotic cycles 13 and 14) (Li et al. 2014). For each peak, we considered a window of 10Kb (at 10bp resolution) and constructed a vector of length 1,000 for each modification. This resulted in a 10x1000 matrix for each Zelda site.

### 3.2. Chromatin-based distance and adjacency functions

To define a distance function between two loci ***x***_***i***_ and ***x***_***j***_, we begin by focusing on one histone modification ***m***:

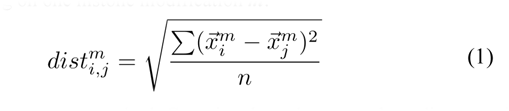

We then use a standard Gaussian kernel to translate distances into weights in the spectral clustering adjacency matrix:

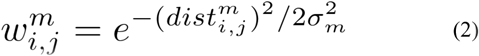

where ***σ***_***m***_ is a modification specific parameter (set below). We then sum the adjacency values over all 10 modifications to obtain one value:

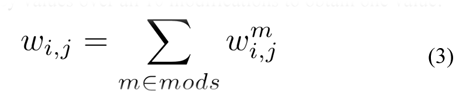

### 3.3. Spectral clustering

Given N <10x1000> matrices **{x**_**i**_**}** that correspond to the histone modification data for each peak, we defined the graph Laplacian:

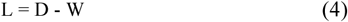

where W is the adjacency matrix (as defined above, Equation 4), and D is the diagonal degree matrix, defined as:

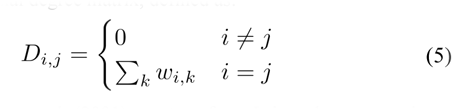

Following Ng et al. (2001), we transform L into the symmetric normalized Laplacian:

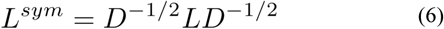

thus normalizing the rows and columns of the graph Laplacian. We then select the first ***k*** eigenvectors of ***L***^***sym***^, project the N data points onto this ***k***-dimensional subspace, and normalize the projected data (rows of length ***k***) to L^2^ norm of 1 (i.e. project each data point to the ***k***-dimensional sphere). Finally, we apply a standard k-means clustering algorithm (where ***k*** here equals the number of eigenvectors ***k*** used for the spectral projection).

To choose ***k***, we applied a standard eigengap (or spectral gap) heuristic, namely calculating the difference between successive eigenvalues (**λ**_**k**_ and **λ**_**k+1**_), and choosing the first to surpass some threshold.

### 3.4. Empirically normalized distance/adjacency function

In Equation 2 we described the Gaussian kernel used to translate average Euclidean distances into adjacency weights. As each genomic track (histone modification, time) presents different absolute values, it is crucial to normalize each Gaussian specifically using its own scaling parameter ***σ***_***m***_. For this, we calculated the empirical distribution of (unnormalized) pairwise distances for each modification ***m*** (Figure 9, purple line), and set ***σ***_***m***_ to be the 10^th^ percentile (red vertical line, Figure 9). This way, we ensured that for each modification exactly 10% of the pairwise distances (Figure 9, purple regions) will have adjacency values larger than exp(−½)=0.606 in the adjacency matrix. This independently normalizes each modification to its own distribution of pairwise distances, thus setting similar “weight” to each modification, allowing for the summation of different adjacency matrices into one (Equation 3).

**Figure 9.**
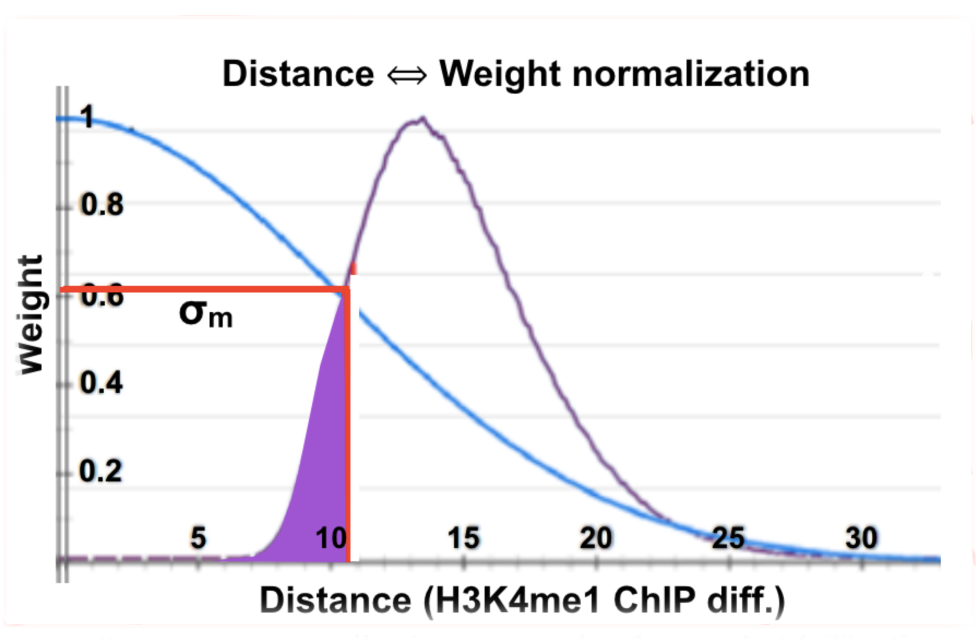
Distance distribution of H3K4me1 at cycle 14. The figure above shows the empirical pairwise distance distribution (purple line) of histone modification H3K4me1 at cycle 14, over pairs of early Zelda peaks <**x**_**i**_**,x**_**j**_>. The vertical red line corresponds to the 10^th^ percentile, in this case distance of 10.36. This value is assigned as ***σ***_***m***_ (horizontal red line). The blue line shows the matching Gaussian kernel function (with σ=10.36), which is used to transform distance (X-axis) to weights (Y-axis).

### 3.5. Genome-wide scanning of similar chromatin loci

Following the spectral clustering of Zelda peak into three classes, we wished to identify novel enhancer regions with similar chromatin signature to that of Zelda cluster 2 peaks. For this, we scanned the genome in (overlapping) 10Kb windows, and cross-correlated the chromatin signature at each window to the average histone modification pattern of cluster 2 (enhancer-like regions). We scored each locus ***i*** by first calculating the Pearson correlation between its surrounding **x**
^
**m**
^
_**i**_ (where ***m*** denotes the histone modification / time point) vs. the chromatin signature for cluster 2 peaks (denoted by ***cl***
^
***m***
^
_***2***_).

This approach, as opposed to just comparing the maximal peak height within each window, gives spatial resolution and identifies the unique shapes we observe for the different modifications (e.g. two symmetric narrow peaks, flaking domains, asymmetrical peak, etc.).

To incorporate prior knowledge and capture the overall complex shape of enhancer-like peaks, namely the typical combination of heights for each modification/time point, we also included a “prior” term ***P***:

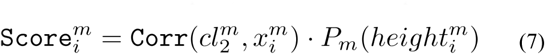

where **P**_**m**_**(height**^**m**^_**i**_) is the empirical probability function of observed peak heights for modification ***m***, and **height**^**m**^_**i**_ denotes the maximal height for modification m in the 10Kb surrounding locus ***i***. We calculated ***P*** using the chromatin signatures around the initial set of early Zelda peaks in cluster 2. This penalizes loci with similar shape (hence, high Pearson correlation coefficients) but in different ChIP magnitude.

Finally, we calculated the overall similarity score for each locus, by summing over the modification specific scores:

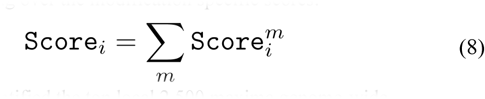

and identified the top local 2,500 maxima genome-wide.

## 4 Discussion

In this work, we developed a computational method aimed to analyze the chromatin landscape surrounding some regions of interest (here, binding of the Zelda protein in early embryonic development of the fruit fly *Drosophila melanogaster*), refined it by clustering into sub-groups of unique characteristics, and then scan the entire genome for novel regions showing similar chromatin characteristics.

As we have shown, much of the methods accuracy stems from the initial re-orientation and clustering procedure, which allows us to derive a specialized mixture-model rather than averaging away the chromatin signal.

We have devised methodologies to simultaneously analyze multiple genomic tracks (i.e. different histones modifications) at different time points, while maintaining the spatial signature of each modification. Such models are more compelling than naive combinatorial models, that only consider the heights of the various modifications but ignore the overall shape. As we have shown, many of the observed signals come in unique shapes, including a dip at the ZLD site with two flanking nucleosomes marked with one modification, all within some domain of different modification.

We have analyzed early Zelda peaks throughout the *Drosophila* genome, and identified three typical chromatin signatures they are associated with, including (1) active promoters, (2) active enhancer regions, and (3) genomic regions that were transiently bound by Zelda. We then characterized each of the classes, and scanned the genome for additional regions matching the enhancer-like chromatin signature. As we have shown, many of the retrieved regions were shown to be bound by either Zelda or GAF, suggesting both act as pioneer factors by making these genomic regions accessible and depositing the appropriate histone modifications, thus marking them for activation during the Maternal-to-Zygotic transition. The entire role of GAF in MZT as well as the possible role for additional factors, is yet to be studied.

While these questions are key to understanding the processes that initiate and shape embryonic gene expression in the fruit fly *Drosophila melanogaster*, the method presented here is more general and would allow – given a query set of input loci – the identification of similar elements in any genome. In that sense, we believe that our method could be of great interest to any genome-wide research projects.

## Acknowledgements

This work has been supported by the Israeli Centers of Excellence (I-CORE) for Gene Regulation in Complex Human Disease (grant no. 41/11) and for Chromatin and RNA in Gene Regulation (grant no. 1796/12), and also by a Marie Cure Integration Grant (no. PCIG13-GA-2013-618327).

